# Thiamin priming to control early blight in potato: investigation of its effectiveness and molecular mechanisms

**DOI:** 10.1101/2024.09.06.611704

**Authors:** Trenton W. Berrian, Matthew L. Fabian, Conner J. Rogan, Jeffrey C. Anderson, Christopher R. Clarke, Aymeric J. Goyer

## Abstract

**Background:** Previous reports in several plant species have shown that thiamin applied on foliage primes plant immunity and is effective in controlling fungal, bacterial, and viral diseases. However, the effectiveness of thiamin against potato (*Solanum tuberosum*) pathogens has seldom been investigated. Additionally, the transcriptomics and metabolomics of immune priming by thiamin have not previously been investigated. Here, we tested the effect of thiamin application against *Alternaria solani,* a necrotrophic fungus that causes early blight disease on potato foliage, and identified associated changes in gene expression and metabolite content.

**Results:** Foliar applications of thiamin reduced lesion size by approximately 33% when applied at an optimal concentration of 10 mM. However, the effect of thiamin on preventing lesion growth was temporally limited, as we observed a reduction of lesion size when leaves were inoculated 4 h, but not 24 h, following thiamin treatment. Additionally, we found that the effect of thiamin on lesion size was restricted to the site of application and was not systemic. Gene expression analysis via RNA-seq showed that thiamin induced the expression of genes involved in the synthesis of salicylic acid (SA) and phenylpropanoids to higher levels than the pathogen alone, as well as fatty acid metabolism genes that may be related to jasmonic acid biosynthesis. Thiamin also delayed the downregulation of photosynthesis-associated genes in plants inoculated with *A. solani*, which is a typical plant response to pathogens, but could also induce a similar repression of primary metabolic pathways in non-infected leaves. Metabolite analyses revealed that thiamin treatment in the absence of pathogen decreased the amounts of several organic compounds involved in the citric acid cycle as well as sugars, sugar alcohols, and amino acids.

**Conclusions:** Our study indicates that thiamin priming of plant defenses may occur through perturbation of primary metabolic pathways and a re-allocation of energy resources towards defense activities.

## Background

Early blight of potato is a disease caused by the fungal pathogen *Alternaria solani* (Family *Pleosporaceae*). The primary symptom of early blight is the presence of necrotic lesions on leaves, often most prevalent in senescing or stressed tissue [1]. Small circular lesions progress into large angular lesions eventually causing localized death. The ensuing reduced photosynthetic ability can result in a dramatic yield reduction [2]. The pathogen can also be symptomatic on tubers, resulting in dry rot symptoms and the formation of dark and sunken lesions on the tuber surface [3], making the tuber unmarketable and unsuited for processing. *A. solani* occurs in nearly all potato growing regions of the world and under many different climates [4]. Environmental conditions such as high moisture and temperature can speed the development of the disease [5, 6]. The conventional approach to controlling early blight in potato is the application of fungicides [7, 8], which result in reduction of disease [9], especially when applied at critical times such as late bulking and tuber maturation [10]. However, many of the major classes of fungicides used to control early blight have been shown to lose effectiveness due to fungicide resistance [8, 11]. Moreover, fungicides can have lasting negative effects on both the environment and human health [12, 13]. Due to environmental and health effects, along with the emergence of fungicide-resistant strains of *A. solani,* more management strategies must be investigated.

Among some of the other common management strategies to prevent *A. solani* outbreaks is the use of resistant cultivars. Unfortunately, breeding for resistant cultivars has been limited due to the multigenic nature of resistance [14]. Biocontrol is another promising route of disease control [15]. However, there is a need for more field-scale research to prove its effectiveness [16]. Additional management strategies include cultural management, such as crop rotation, the removal of alternate hosts such as *S. nigrum* and *S. carolinense,* the utilization of disease-free seed, and irrigation practices [15, 17, 18]. Although cultural management practices are effective at partially controlling the disease, full control is often unattainable. Typically, a mixture of fungicide applications, resistant varieties, and cultural controls are needed to mitigate early blight disease. Accordingly, there is a considerable need to identify more effective tools to prevent yield losses resulting from early blight disease.

An alternative strategy for managing plant disease is the application of chemical elicitors for immune priming of plant defenses, characterized by the activation of host plant defense responses, such as transcription of defense genes and the synthesis of phytoalexins, in advance of pathogen infection. Amongst chemical elicitors, B group vitamins have recently received attention as natural plant products that are able to prime plant defenses and reduce disease incidence [19]. Thiamin (vitamin B1), in its pyrophosphorylated form thiamin diphosphate (ThDP), is a cofactor for key enzymes of carbohydrate, amino acid, and fatty acid metabolism [20, 21]. In particular, ThDP is a cofactor for both mitochondrial and chloroplastic pyruvate dehydrogenases, which are involved in glycolysis and *de novo* fatty acid biosynthesis, respectively. ThDP is also a cofactor of transketolase, a key enzyme of the oxidative pentose phosphate pathway and the Calvin cycle. Another key enzyme of primary metabolism that uses ThDP as a cofactor is 2-oxoglutarate dehydrogenase, which has essential roles in the citric/tricarboxylic acid (TCA) cycle, nitrogen assimilation, and amino acid metabolism. ThDP is essential for the synthesis of the branched-chain amino acids valine, leucine, and isoleucine as a cofactor for acetolactate synthase, and also serves as a cofactor for 1-deoxy-D-xylulose-5- phosphate (DXP) synthase, which synthesizes DXP, a precursor of isoprenoids via the mevalonate-independent pathway.

Thiamin has also been shown to prime plant defenses when externally applied to plant foliage in advance of pathogen challenge [22–25]. Priming of plant defenses with thiamin was demonstrated to slow or stop infections from fungal, viral, and bacterial pathogens in a variety of hosts, including *Arabidopsis*, soybean, rice, grape, tobacco and cucumber [22, 23, 26, 27]. Plant defenses triggered by thiamin application include callose deposition, phytoalexin production, pathogenesis-related (PR) gene expression, and production of reactive oxygen species (ROS) [19]. An increase in biosynthesis of secondary metabolites such as terpenoids, phenylpropanoids and antioxidants was found in grapevine upon treatment with thiamin, and the production of such molecules are most likely regulated through molecules such as lipoxygenases, which are also upregulated upon thiamin treatment on grapevine [26]. These molecular changes involve the SA-dependent signaling pathway in *Arabidopsis* [23]. In potato, application of thiamin decreased the viral titer of potato virus Y [28]. However, no other studies have been conducted to address the potential of thiamin as a priming agent in potato against other pathogens. It is also unclear how thiamin primes plant defenses in any of the plant pathosystems studied so far. Possible mechanisms include thiamin functioning as an enzymatic cofactor, and possible subsequent metabolic reorganization, or through a non-cofactor, yet-to-be-identified role.

In this study, our first objective was to evaluate the effectiveness of thiamin priming treatments against foliar *A. solani* infections in potato. Second, after demonstrating that thiamin treatment decreased symptoms caused by *A. solani*, we characterized the molecular mechanisms of thiamin priming in potato by analyzing changes of gene expression by RNA-seq and changes in metabolites by gas chromatography mass spectrometry (GC-MS).

## Methods

### Plant growth

All experiments were done with the potato variety Russet Norkotah, and in one experiment as noted in the text below, Russet Burbank was used as well. Both varieties were chosen because of their susceptibility to early blight. Plantlets were grown for three to four weeks on solid Murashige and Skoog (MS) medium (4.6 g l^-1^ MS-modified BC potato salts, 30 g L^-1^ sucrose, 100 mg L^-1^ myo-inositol, 2 mg L^-1^ glycine, 0.5 mg L^-1^ nicotinic acid, 0.5 mg L^-1^ pyridoxine, 0.1 mg L^-1^ thiamine, pH 5.6) before being transferred to pots containing a mixture of sand and potting soil (Sunshine Mix #4) (v/v 1:4) and slow-release fertilizer (Osmocote Plus). Plants were then grown in greenhouses for an additional 5-7 weeks before utilization in experiments. Plants received 14 h daily light exposure, and supplementary lighting was provided with 400 W high-pressure sodium lamps. Greenhouse temperature was maintained at 21°C day and 18°C night. For whole plant assays, plants were placed in a 1.22 m x 0.91 m x 2.44 m humidity chamber made from PVC piping and clear plastic.

### Thiamin foliar treatments

Thiamin (Millipore Sigma, Catalog No W332208, ≥98%) solutions were prepared in deionized water that included Tween 20 at 250 µg L^-1^ to facilitate dispersion of thiamin to the foliage; mock solution consisted of Tween 20 at 250 µg L^-1^ in deionized water. Treatment solutions were sprayed onto foliage via a handheld spray bottle until runoff (approximately 30 mL per plant). Thiamin was applied at the given concentrations (0, 1, 5, 10, 25, 50 mM) and timepoints (4 h, 28 h) prior to pathogen inoculation as noted.

### *A*. *solani* inoculations

*Alternaria solani* strain BMP 183 [29] was grown on V8 agar medium plates (10% clarified V8 juice, 1.5 % CaCO3, and 12.7% Agar) from a glycerol stock kept at -80°C. After three days growth, the pathogen was sub-cultured onto fresh V8 agar plates and grown under continuous light until complete coverage of the plate was observed (15-21 days). Plates were then covered with 5 mL deionized water and conidia were gently dislodged using a plastic spreader. Conidia were then transferred into a 50-mL falcon tube, vortexed to release spores from mycelium, and filtered through four layers of cheesecloth. Conidia were counted via hemocytometer and concentration was adjusted to 15,000-30,000 spores mL^-1^ with deionized water for all assays except systemic tests where spore concentration was adjusted to ∼6,000 spores mL^-1^. Spore concentration used for each experiment is indicated in the legends of figures and table.

### Detached leaflet inoculations

After mock or thiamin treatment on whole plants, four leaflets were removed using a sterile scalpel from the 3^rd^ and 4^th^ leaves from the top of each plant. Leaflets were rinsed with deionized water to remove any treatment residue and thiamin precipitate and allowed to dry before inoculations. Laboratory wipes (Kimwipe) (2.5 x 2.5 cm) were wrapped around the petiole of the leaflet. Leaflets were then arranged on pipette tip holders in 1020 garden trays (Greenhouse Megastore) and separated by treatment group, with one biological replicate per tray; four biological replicates per group were used in all assays. The detached leaflets were then drop-inoculated with four equally spaced drops of 20 µL inocula on the adaxial side of the leaflet. Three leaflets per plant were used for *A. solani* inoculation, and one inoculated with deionized water only as control. After pathogen inoculation, wipe squares were saturated daily with 200 µL deionized water to keep leaflets hydrated. To maintain high humidity for optimum infection, 250 mL of reverse osmosis water was added to the bottom of the trays, which were covered with clear plastic domes and sealed with packing tape. Trays were placed in a dark growth chamber immediately post-inoculation for 14 h at 22 , then under a 10 h photoperiod until lesions were large enough for measurement, at 3-4 days post-inoculation. To evaluate the efficacy of thiamin as a priming agent for systemic immunity against *A. solani*, several leaves of each plant were covered using plastic zip-lock bags to protect them from thiamin treatments performed as described above. Four hours after thiamin treatment, we collected unbagged and bagged leaflets for a detached leaf assay.

### Whole plant inoculations

Plants were placed in the humidity chamber, and leaflets were inoculated directly on the plant with two to six 10 µL drops of inocula per leaflet, depending on leaflet size. In collecting leaf disks for RNA-seq (see below), leaflets all had six drops of inocula. Mock-inoculated and *A. solani*-inoculated samples were from different leaves of the same plant in both mock- and thiamin-treated plants across all timepoints. Four biological replicates (one biological replicate = one plant) were used for each treatment, with a minimum of four leaves per plant. To provide moisture (relative humidity > 90%) and encourage disease development, a humidifier (AquaOasis) was used for 2 h post-inoculation in the late afternoon (16:00) and subsequently for 2 h every morning and evening. A black shade tarp was draped over the chamber immediately post-inoculation in the late afternoon and removed the following morning.

### Lesion measurements

Disease severity was determined after three to four days by modeling the lesions using the trace function and measuring the area of each lesion in mm^2^ via ImageJ [30]. For the detached leaf assay, the value recorded for each plant is the mean lesion area across all 12 lesions. For the whole plant assay, all lesions on a single plant were measured via ImageJ and reported as a single mean lesion area per biological replicate. For the systemic resistance assay, lesion diameter was recorded instead of area.

### RNA extraction

Whole plants (n=3) were spray treated with 10 mM thiamin or mock solution. Four hours post-treatment, three leaflets from the same leaf were inoculated with *A. solani*. Four biological replicate samples (leaves from one plant = one biological replicate) were collected from both *Alternaria-*inoculated and non-inoculated leaves on both thiamin-treated and mock-treated plants at three time points (12, 24, and 48 hours post-inoculation (hpi)). For each time point, treatment groups were assigned as follows: “mock_Alt_local”, mock-treated, *A. solani*-inoculated leaves; “mock_distal”, mock-treated, non-inoculated/distal leaves; “thi_Alt_local”, thiamin-treated, *A. solani*-inoculated leaves; and “thi_distal”, thiamin-treated, non-inoculated/distal leaves (Fig. S1). For samples from each time point and treatment group, an 8-mm diameter hole punch was used to take six leaf discs from the inoculated area, weighing a total of 50-60 mg per sample. Non- inoculated, mock- and thiamin-treated samples were collected concurrently with inoculated samples and were therefore assigned hpi designations of 12, 24, and 48 hpi accordingly. Samples were immediately frozen in liquid nitrogen and stored at -80°C until RNA extraction. Frozen leaf tissue was first homogenized using a mortar and pestle, and total RNAs were extracted using the PureLink^TM^ RNA Mini Kit (Invitrogen) using the manufacturer’s instructions. Trace genomic DNA was removed via DNase I treatment via DNA-*free*^TM^ Kit (Invitrogen). RNA was precipitated via one volume of 4 M lithium chloride incubated at 4°C overnight. After centrifugation at 13,250 g for 30 min at 4°C, the pellet was washed with 200 µL 70% ethanol and resuspended in RNase-free water. RNA integrity was evaluated via gel electrophoresis, potential contamination was assessed via a NanoDrop One^C^ (ThermoScientific) spectrophotometer, using absorbance ratios of A260/280 and A260/230 ≥ 2.0 for cut-off. RIN values were obtained via Agilent Bioanalyzer 2100 and ranged in value from 6.6 to 9.

### qRT-PCR

RNA concentrations and purities were analyzed via a DeNovix DS-11 spectrophotometer (DeNovix Inc., Wilmington DE, USA). Synthesis of cDNA was conducted via ProtoScript II First Strand cDNA Synthesis Kit (New England Biolabs, Ipswich MA, USA), using manufacturer’s instructions for reverse transcription of total RNA. cDNA samples were diluted 100-fold for qRT-PCR. Primers for qRT-PCR were designed from target mRNA sequences, spanning introns where possible, and synthesized by Integrated DNA Technologies, Inc. (Coralville IA, USA). Primer sequences are as follows: StGAPDH (Fwd: GCTCATTTGAAGGGTGGTGC, Rev: AGGGAGCAAGGCAATTTGTG); StPR1 (Fwd: AATGTGCAAGCGGACAAGTG, Rev: TCCGACCCAGTTTCCAACAG). qRT-PCR was carried out via an iProof SYBR Green Supermix kit and CFX96 thermocycler (Bio-Rad Laboratories, Hercules CA, USA) per manufacturer’s instructions. Two technical replicates were included for each of three biological replicates per treatment group and time point. Relative expression (RE) was calculated via the 2^-ΔΔCt^ method [31], with the lowest-expression samples for each time point designated as the calibrator.

### RNA-sequencing

Total RNA was sent to Novogene (Sacramento, CA) for sequencing via Illumina platform. Messenger RNAs were first purified from total RNA using poly-T oligo-attached magnetic beads. Random hexamer primers were used for first strand cDNA synthesis followed by a second strand cDNA synthesis via Illumina NovaSeq platform. After end repair, A-tailing, adapter ligation, size selection, amplification, and purification, the 156-bp paired-end libraries were sequenced using the Illumina NovaSeq 6000 Sequencing System. Adapters used for paired-end sequencing were as follows: 5’ Adapter: 5’- AGATCGGAAGAGCGTCGTGTAGGGAAAGAGTGTAGATCTC- GGTGGTCGCCGTATCATT-3’; 3’ Adapter: 5’-GATCGGAAGAGCACACGTCTGAAC- TCCAGTCACGGATGACTATCTCGTATGCCGTCTTCTGCTTG-3’.

### RNA-seq data analysis

FASTQ file read quality was evaluated via FastQC [32], and adapters were trimmed via Trim Galore! (https://www.bioinformatics.babraham.ac.uk/projects/trim_galore/) with default options, specific adaptor sequences, and filtering for Phred score ≥ 20 [33]. Genome indexing and mapping of trimmed, filtered reads to the potato reference genome DM_1-3_516_R44 v6.1 (http://spuddb.uga.edu/dm_v6_1_download.shtml) was performed via HISAT2 (https://daehwankimlab.github.io/hisat2/) with default options. Output SAM files were converted to BAM format and mapping quality was analyzed via Samtools (http://www.htslib.org/). The High Confidence Gene Model Set annotations file for DM v6.1, in GFF3 format, was converted to GTF format via gffread (http://ccb.jhu.edu/software/stringtie/gff.shtml). Reads were aligned to annotated gene models from the high-confidence gene model GTF via featureCounts (http://subread.sourceforge.net/) with the following options: -p; --countReadpairs; and – transcript.

The DESeq2 package [34] in R (version 1.34.0) was used to generate lists of differentially expressed genes (DEGs) for each comparison group. Genes were differentially expressed between groups only if adjusted *p* ≤ 0.05 and |log_2_(fold-change)| ≥ 2.0. The comparison groups were assigned as follows: “response to Alt (local)”, mock_Alt_local vs. mock_distal; “response to Alt (thi)”, thi_Alt_local vs. thi_distal; “response to thi”, thi_distal vs. mock_distal; and “response to thi (Alt), thi_Alt_local vs. mock_Alt_local (Fig S1). Gene Ontology (GO) term assignments (DM_1-3_516_R44_potato.v6.1.working_models.go.txt.gz) for the annotated genes in the DM v6.1 genome were downloaded from http://spuddb.uga.edu. GO term enrichments for DEGs were identified via the clusterProfiler (https://github.com/YuLab-SMU/clusterProfiler) package in R version 4.1.2. “biological process”, “molecular function”, and “cellular component” terms were analyzed for each list of upregulated and downregulated DEGs, respectively, compiled from the three comparisons and time points and the “universe” option set to the population of genes with ≥ 1 read across all FASTQ files.

### Metabolite analysis by GC-MS

Leaf metabolites collected at 1, 6, and 12 h after thiamin treatment were analyzed by GC-MS as previously described [35]. The difference in sampling times between RNA-seq and metabolite analysis is explained in the text below. Briefly, full leaflets were selected from three plants (one plant = one biological replicate, each plant had every timepoints) sprayed with 10 mM thiamin or mock solution at 1, 6, and 12 hours post treatment (hpt) and frozen in liquid nitrogen. Three biological replicate samples per treatment group for each time point, each of approximately 50 mg mass, were added to 700 µL extraction solvent (water:methanol:chloroform (1:2.5:1)) with an internal standard (40 µg/ml ribitol). Samples were placed on ice for 5 minutes on a shaking platform rotating at 130 rpm, and then centrifuged at 4***°***C for 2 minutes at 21,000 x *g* to pellet cellular debris. The supernatant was transferred to a clean microcentrifuge tube and 280 µL of water was added to separate the aqueous phase from the organic phase. After a 2-minute centrifugation at 21,000 x *g*, the upper aqueous phase was collected and placed into a clean microcentrifuge tube. The samples were frozen at -80 ***°***C, then placed into a centrifugal vacuum concentrator and lyophilized to dryness overnight. Dried samples were stored at -80***°***C until further analysis. A no tissue extraction control (*i.e*., reagent blank) was included to assess if detected peaks are plant tissue-specific. Dried samples were resuspended in 20 µL 30 mg/mL methoxyamine hydrochloride in pyridine and incubated at 37°C for 1.5 h, with vigorous shaking. Next, 40 µL of N-methyl-N-(trimethylsilyl) trifluoroacetamide with 1% trimethylchlorosilane was added and the samples were incubated at 37°C for 30 additional minutes with vigorous shaking. Metabolites were separated in an Agilent 7890B GC system and detected with an Agilent 5977B MSD in EI mode scanning from 50 m/z to 600 m/z. Mass spectrum analysis, component identification and peak area quantification were performed with AMDIS [36]. Fold-change and pathway enrichment analyses were performed via MetaboAnalyst 6.0 (https://www.metaboanalyst.ca/MetaboAnalyst/), with independent comparisons between thiamin- and mock-treated samples at each time point. Metabolites with at least two-fold difference between treatment groups, and with raw p-value < 0.05 were selected for pathway enrichment analysis via hypergeometric test and the *Arabidopsis thaliana*-derived KEGG pathways. Pathway enrichment p-values were manually adjusted via the post-hoc Benjamini-Hochberg method to control the false discovery rate [37].

### Data reporting and statistical analyses

Statistical analysis of assays with multiple comparisons was performed via one-way ANOVA followed by a Tukey’s HSD test for multiple comparisons. Assays incorporating single comparisons were analyzed via Student’s *t*-test or Welch’s *t*-test. RNA-seq and GO enrichment analyses incorporated the post-hoc Benjamini-Hochberg method to control the false discovery rate. Unless otherwise stated, statistical analyses and plots were produced using Microsoft Excel, Venny [38] and/or R version 4.1.2.

## Results

### Thiamin reduces early blight severity in a dose- and time-dependent fashion

To determine the optimal dosage of thiamin for foliar applications, we tested the effect of five different concentrations of thiamin against *A. solani* lesion size via a detached leaf assay. In two independent trials, leaflets from plants treated with 1, 5, 10, 25, and 50 mM thiamin and mock solution were removed and inoculated with *A. solani*. Early blight lesion area was attenuated at all dosage levels of thiamin, in comparison to mock-treated leaves, however statistical significance in lesion size was observed only for the 10 mM dosage (32 and 52% decrease in lesion size in trials 1 and 2, respectively) (Table 1). For this reason, a dosage of 10 mM was used for all subsequent experiments.

**Table 1.** Average lesion area of *A. solani* on plants treated with increasing concentrations of thiamin in two independent trials. Data are means ± S.D. of four biological replicates. Identical letters indicate no statistical difference between treatments as determined by ANOVA and Tukey’s test. Spore concentrations were 21,641 and 27,188 spores per mL in trial 1 and 2, respectively.

| Thiamin<br>concentration<br>(mM) | Lesion size (mean ± SD) | |
| --- | --- | --- |
|  | Trial 1 | Trial 2 |
| 0 | 41.4 ± 8.2 <sup>a</sup> | 52.6 ± 16.6 <sup>a</sup> |
| 1 | 31.9 ± 4.4 <sup>ab</sup> | 34.2 ± 17.0 <sup>ab</sup> |
| 5 | 29.3 ± 4.5 <sup>ab</sup> | 34.6 ± 8.9 <sup>ab</sup> |
| 10 | 28.3 ± 4.0 <sup>b</sup> | 25.2 ± 1.6 <sup>b</sup> |
| 25 | 30.6 ± 6.2 <sup>ab</sup> | 28.7 ± 5.5 <sup>ab</sup> |
| 50 | 31.4 ± 3.8 <sup>ab</sup> | 30.1 ± 5.4 <sup>ab</sup> |

The durability of thiamin treatment in the reduction of *A. solani* lesion size was tested via a detached leaf assay in which *A. solani* inoculations were performed at 4 vs. 28 h post-thiamin treatment in two trials (Fig. 1A). While thiamin was effective in reducing lesion size when leaflets were inoculated with *A. solani* at 4 hpt, there was no observable effect at 28 hpt, indicating that the effect of thiamin application is transitory. Consequently, all following thiamin treatments were performed at the optimal 10 mM concentration and with inoculations occurring at 4 hpt.

**Fig 1.**
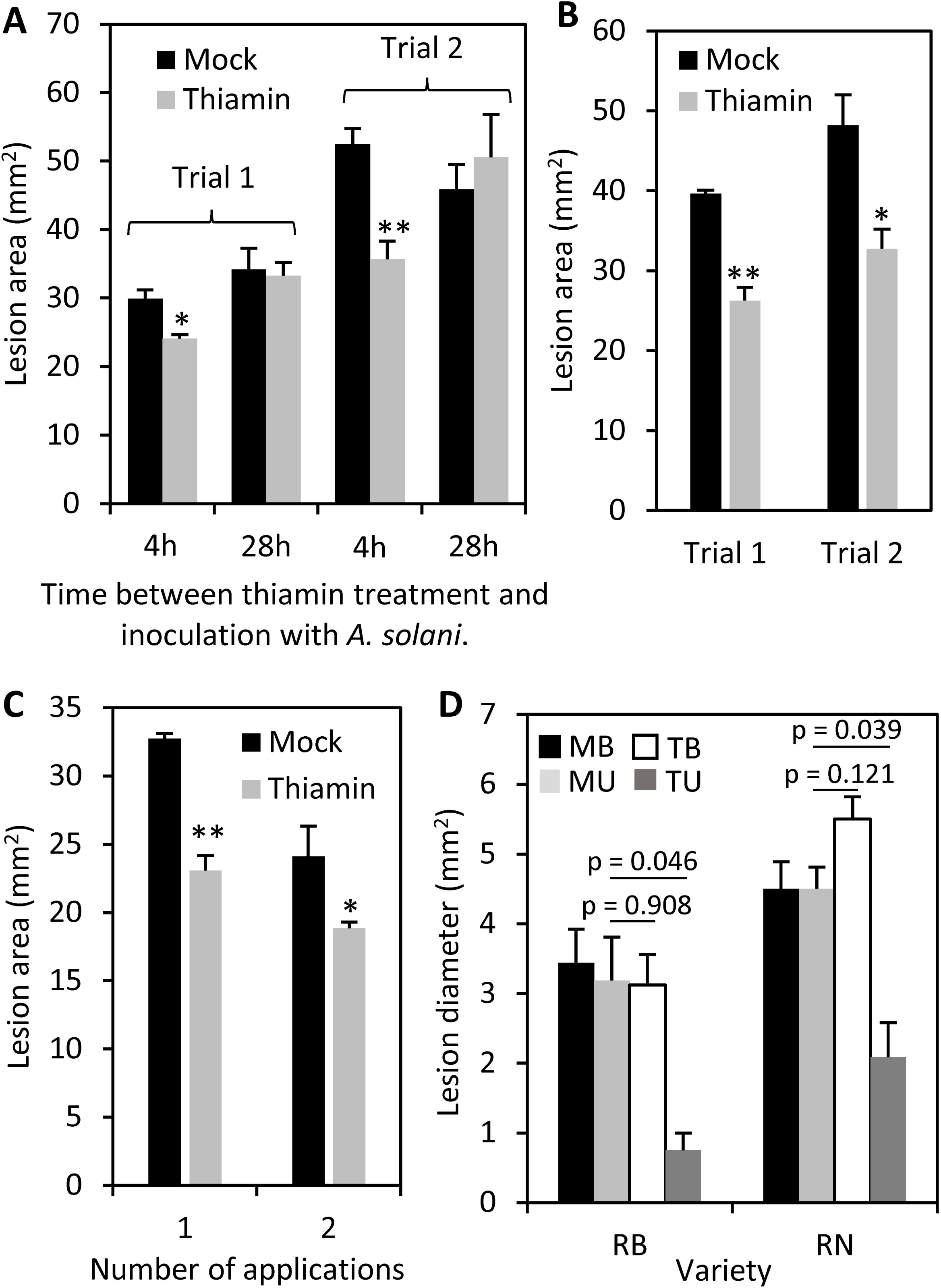
Effects of thiamin foliar application on lesions caused by *A. solani*. (**A**) Average lesion area of *A. solani* on plants treated with either 10 mM thiamin or a mock solution and inoculated either 4 or 28 h after treatment in two independent trials. Spore concentrations (spores per mL) were 21,563 (4 h) and 24,688 (24 h) for trial 1, and 16,785 (4 h) and 18,125 (24 h) for trial 2. (**B**) Average lesion area of *A. solani* on whole plants treated with either 10 mM thiamin or mock solution and inoculated 4 hr after treatment in two independent trials. Spore concentrations (spores per mL) were 15,469 and 17,813 for trial 1 and trial 2, respectively. (**C**) Average lesion area of *A. solani* in plants treated with one or two thiamin foliar applications. The second thiamin application was done seven days after the first one on the same plants. Spore concentrations (spores per mL) were 15,468 and 21,562 for trial 1 and trial 2, respectively. (**D**) Average lesion area of *A. solani* on bagged and unbagged leaves from plants treated with thiamin or a mock solution as determined by detached leaf assay. Spore concentration (spores per mL) was 5,875. Treatments: MB = <u>B</u>agged leaves from <u>m</u>ock plants, MU = <u>U</u>nbagged leaves from mock plants, TB = <u>B</u>agged leaves from thiamin-treated plants, TU = <u>U</u>nbagged leaves from thiamin-treated plants. Varieties: RB = Russet Burbank, RN = Russet Norkotah. All data are means ± S.E. of four biological repetitions, except (**D**) which had three and six biological replicates for mock- and thiamin-treated plants, respectively. Asterisks indicate a p-value of ≤ 0.05 (*) or ≤ 0.01 (**) when compared to mock treatment as determined by Student t-test, except for (**D**) where Welch’s T test was used for comparing MU vs. TU and MU vs. TB.

Utilizing the optimized thiamin dosage and timing, we surveyed the efficacy of thiamin in priming immunity against *A. solani* in whole plants. We observed that thiamin reduced the size of lesions in two additional independent trials by 34% and 32%, respectively (Fig. 1B). To explore whether repeated thiamin applications desensitize potato to the corresponding immunity priming, we incorporated a detached leaf assay utilizing multiple applications of thiamin. Whole plants were sprayed with 10 mM thiamin or mock solution, and detached leaflets were inoculated with *A. solani* 4 hpt. After 7 additional days, plants were sprayed with a second application of thiamin, and a second set of detached leaflets, that were also previously treated with thiamin or mock solutions, were removed and inoculated with *A. solani*. Across the first and second applications, thiamin reduced lesion size by 29% and 22%, respectively, suggesting that repeated thiamin applications at the tested time points do not desensitize the plants to thiamin priming (Fig. 1C).

To evaluate the efficacy of thiamin as a priming agent for systemic immunity against *A. solani*, we designed a bagged leaf assay incorporating two cultivars of potato. Whole plants were treated with thiamin with multiple leaves enclosed in plastic bags, to shield those leaves from direct contact with thiamin. Bagged and unbagged leaves were removed 4 hpt and incorporated in a detached leaf inoculation assay, from which we observed that unbagged, thiamin-treated leaves had reduced lesion size in comparison to bagged leaves (Fig 1D). Lesions of bagged, thiamin-treated leaves were similar to those of unbagged and bagged mock-treated leaves suggesting that the thiamin treatments only locally reduce *A. solani* lesion development.

### RNA-seq experimental design

To shed light on potential regulatory and response pathways of thiamin priming against *A. solani*, we utilized RNA-seq to investigate the transcriptome. Because we observed thiamin only acts locally to attenuate early blight disease symptoms, the RNA-seq experiment was designed to identify genes specifically involved in local response to *A. solani* infection and thiamin treatments. Namely, locally inoculated *A. solani* leaves were compared against non-inoculated leaves from *A. solani*-inoculated plants (on distal leaves) to reduce the population of DEGs to only plant genes involved in local, but not systemic, response to *A. solani.* This design also replicates possible field conditions for early blight disease pressure, in which individual plants could possess a mix of both locally-infected and distal, non-infected foliage. Plants were separately treated with thiamin or mock solution, and at 4 hpt, leaves were inoculated with either *A. solani* (“local”) or a mock solution (“distal”), after which leaf samples were collected 12, 24, and 48 hpi; these time points correspond to no visible lesion, beginning of lesion, and clear lesion, respectively (Fig. S2). Furthermore, a qRT-PCR assay incorporating thiamin treatment and *A. solani* infection indicated that expression of the defense marker gene *PR-1* was induced by *A. solani* infection, in both mock- and thiamin-treated leaves, at 24 and 48 hpi, and that thiamin significantly induced *PR-1* in non-inoculated leaves at 24 hpi (Fig. S3). For each time point, samples were collected in four treatment groups: mock-treated and inoculated (“mock_Alt_local”); mock-treated and non-inoculated (“mock_distal”); thiamin-treated and inoculated (“thi_Alt_local”); and thiamin-treated and non-inoculated (“thi_distal”) (Fig. S1).

Across all samples, 88.14% of trimmed reads mapped to the potato reference genome, and 86.67% of paired reads aligned to annotated gene models (Table S1). Principal Component Analysis was conducted on paired read counts aligned to genome features and revealed strong concordance among the biological replicates (Fig. S4). Differentially expressed genes (DEGs) were computed for four treatment comparisons (Fig. S1). To survey the local transcriptional response to *A. solani* (“response to Alt (local)”), mock-treated, locally inoculated (“mock_Alt_local”) samples were compared to samples derived from mock-treated, distal/non-inoculated tissue (“mock_distal”). The response to *A. solani* was also surveyed in the context of thiamin-treated plants (“response to Alt (thi)”), via comparison of “thi_Alt_local” to “thi_distal”. Direct response to thiamin (“response to thi”) was assessed using samples from distal/non-inoculated tissue: “thi_distal” vs. “mock_distal”. The response to thiamin in the context of an *A. solani* infection (“response to thi (Alt)”) was also surveyed in the background of locally inoculated tissue (“thi_Alt_local” vs. mock_Alt_local”), however only 16 DEGs were identified at 12 hpi, 139 DEGs were identified at 24 hpi, and no DEGs were identified at 48 hpi, suggesting that transcriptional response to *A. solani* largely masks the transcriptional response to thiamin. Distributions of DEG expression fold-change and statistical significance by comparison group were visualized via volcano plots (Fig. S5).

### Overall transcriptional responses to *A. solani* and thiamin

In the “response to thi” comparison (thiamin-treated vs. non-treated samples from distal/non-inoculated leaves), comparatively few DEGs were identified, including 111, 92, and 168 DEGs at 12, 24, and 48 hpi, respectively, with two DEGs shared across all timepoints (Fig. 2A; Tables S2-S4). The response to thiamin was also surveyed in the context of *A. solani* infection (“response to thi (Alt)”), yielding 16 DEGs at 12 hpi and at 24 hpi, 139 DEGs, all downregulated (Tables S5-S6). In “response to Alt (local)” (locally inoculated vs. distal/non-inoculated leaves; mock treatment), there were 2,421, 2,258, and 3,836 DEGs (Fig 2B; Tables S7-S9), compared to 1,217, 2,576, and 2,824 in “response to Alt (thi)” (locally inoculated vs. distal/non-inoculated leaves; thiamin treatment), at 12, 24, and 48 hpi, respectively (Fig 2C; Tables S10-S12). A total of 1,175 DEGs were common to all 3 timepoints in “response to Alt (local)”, in comparison to 715 DEGs common to all time points in “response to Alt (thi)” (Fig. 2B-C). In a comparison of the “response to Alt (local)” and “response to Alt (thi)” transcriptional responses across time points, the former exhibited a larger volume of DEGs at 12 and 48 hpi, with the inverse observed at 24 hpi (Fig. 2D-F). However, DEGs exhibited a large degree of overlap between both groups, with 954, 1,508, and 2,389 DEGs identified in both groups at 12, 24, and 48 hpi, respectively (Fig. 2D-F). Taken together, these results suggest a much more robust transcriptional response to *A. solani* infection than thiamin treatments and a shift in transcriptional response to *A. solani* in the context of thiamin pretreatment.

**Fig 2.**
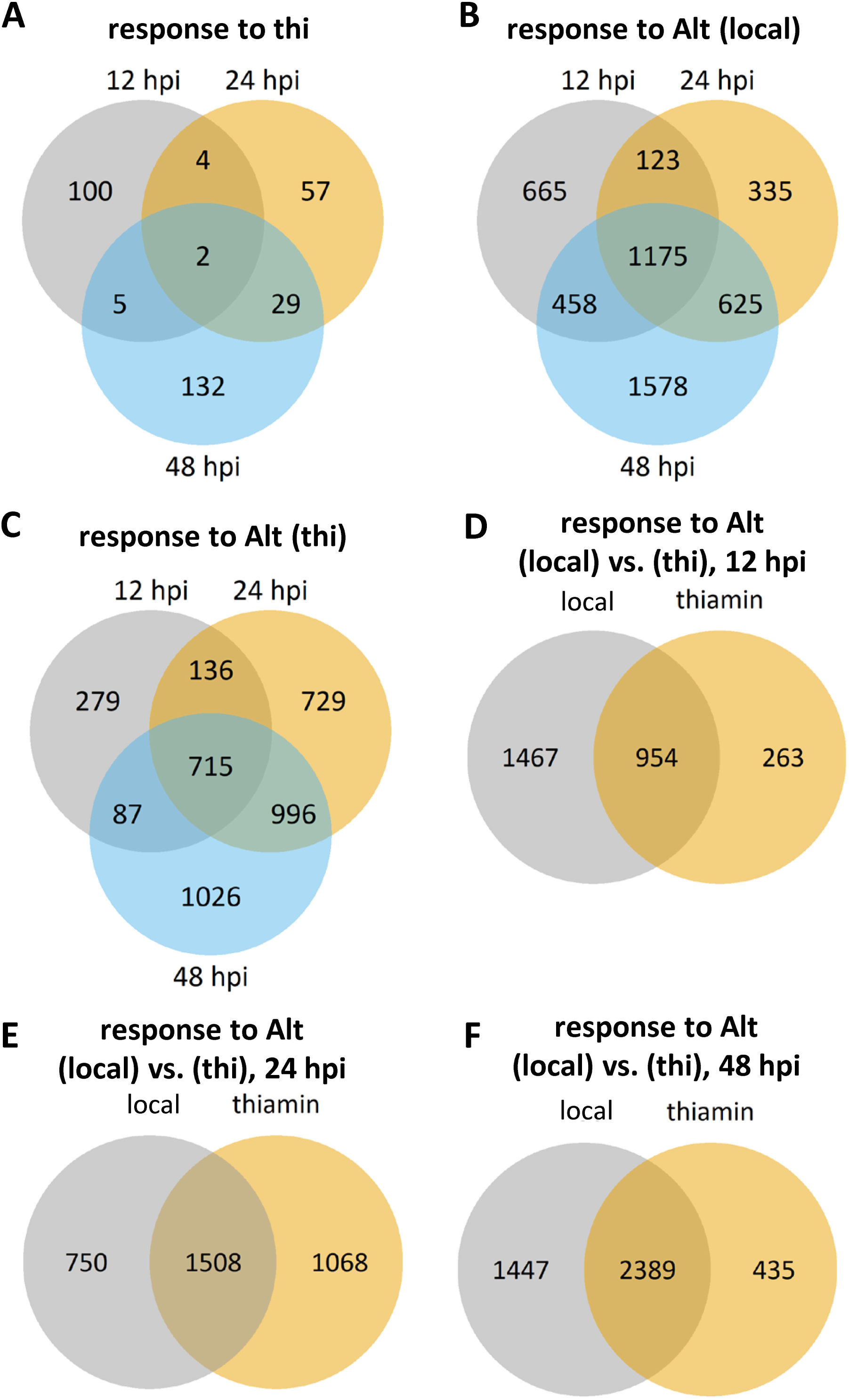
Comparison of differentially expressed genes (DEGs) between treatment groups and timepoints. (**A**) Number of DEGs in “response to thi” (thi_distal vs. mock_distal) at 12, 24, and 48 hpi. (**B**) Number of DEGs in “response to Alt (local)” (mock_Alt_local vs. mock_distal) at 12, 24, and 48 hpi. (**C**) Number of DEGs in “response to Alt (thi)” (thi_Alt_local vs. thi_distal) at 12, 24, and 48 hpi. (**D-F**) Comparisons of DEGs between mock- and thiamin-treated plants in response to *A. solani* (“response to Alt (local)” and “response to Alt (thi)”) at 12 (**D**), 24 (**E**), and 48 (**F**) hpi.

### Thiamin treatment influences transcriptomic and metabolomic pathways in primary metabolism

To probe potential pathways through which thiamin primes immune response, using DEGs in the direct response to thiamin, absent pathogen (“response to thi”; Tables S2-S4), we performed Gene Ontology (GO) enrichment analysis on upregulated and downregulated DEGs for each of the three time points. At 24 hpi, upregulation of two fatty acid desaturase genes contributed to significant enrichment for multiple fatty acid-associated GO terms, and upregulated glycosyl hydrolase/chitinase genes conferred enrichment for terms corresponding to chitinases (Fig. 3A; Tables S3, S13). At 48 hpi, enriched GO terms for upregulated DEGs corresponded to protease inhibitor and cytochrome P450 genes, as well as two peroxidases associated with GO terms for fatty acid alpha-oxidation and responses to ROS, SA, and nitric oxide (NO). For downregulated genes, significantly enriched GO terms were observed only at 12 hpi and were broadly associated with photosynthesis GO terms due to a group of underlying genes encoding chlorophyll-binding proteins (Fig. 3B; Tables S2, S13). We also analyzed the transcriptomic response to thiamin in the context of infection (“response to thi (Alt)”), using leaf tissue locally infected with *A. solani*. A variety of significantly enriched GO terms were identified from upregulated DEGs at 12 hpi, with most corresponding to a single DEG (Fig. 4A; Tables S5, S14). Downregulated DEGs conferred significant enrichment of GO terms at 12 and 24 hpi, with a number of terms associated with the chloroplast, as well as photosynthesis and Calvin cycle processes (Fig. 4B; Tables S5-S6, S14). These results suggest a comparatively limited transcriptional response to thiamin treatment that is at least partially masked/obscured by *A. solani* infection at the tested time points.

**Fig 3.**
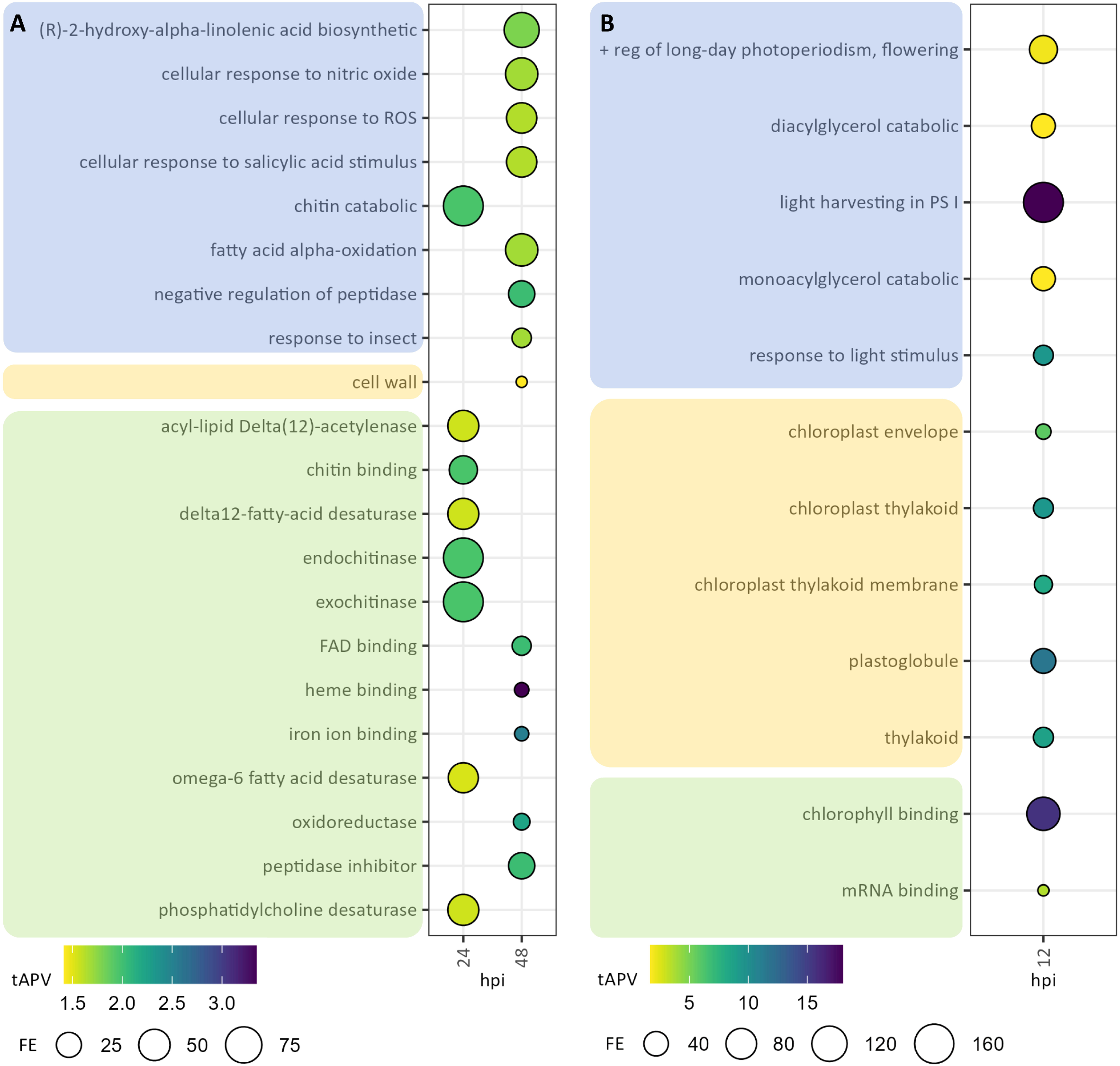
GO term enrichment of upregulated (A) and downregulated (B) DEGs in “response to thi”. DEGs were selected on the basis of |log_2_(fold-change)| ≥ 2 and adjusted p-value < 0.05. Enriched GO terms were selected on the basis of adjusted p-value (APV) < 0.05. [column rows]. “hpi”, hours post-inoculation. “tAPV”, -log_10_-transformed APV. “FE”, fold-enrichment. Blue- shaded terms, “Biological Process”; orange-shaded terms, “Cellular Component”; green-shaded terms, “Molecular Function”.

**Fig 4.**
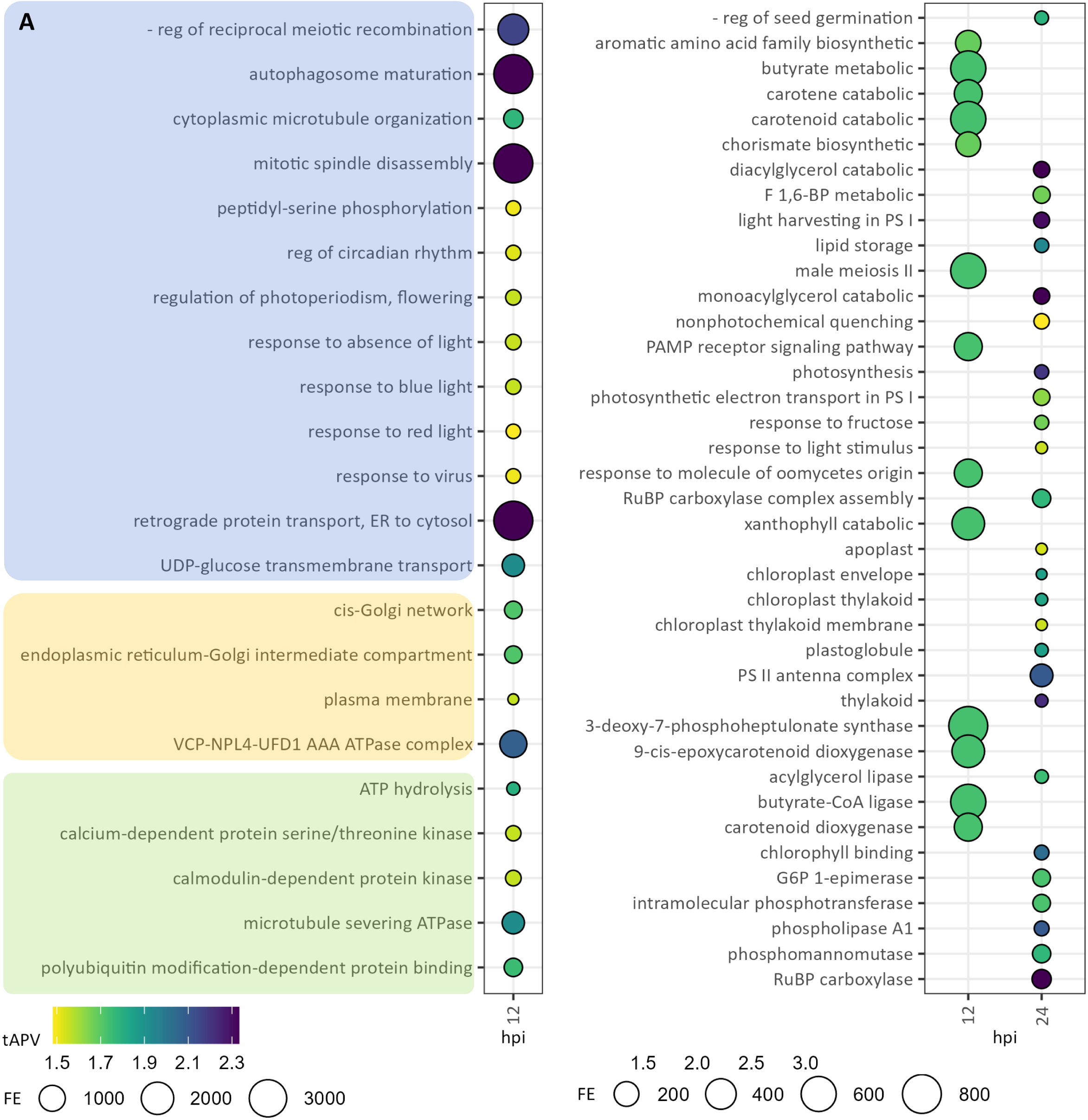
GO term enrichment of upregulated (A) and downregulated (B) DEGs in “response to thi (Alt)”. DEGs were selected on the basis of |log2(fold-change)| > 2 and adjusted p-value < 0.05. Enriched GO terms were selected on the basis of adjusted p-value (APV) < 0.05. “hpi”, hours post-inoculation. “tAPV”, -log10-transformed APV. “FE”, fold-enrichment. Blue-shaded terms, “Biological Process”; orange-shaded terms, “Cellular Component”; green-shaded terms, “Molecular Function”.

Accumulation of ThDP, the cofactor form of thiamin, via mutation in the ThDP riboswitch, has been shown to increase the activities of the thiamin-dependent enzymes pyruvate dehydrogenase, α-ketoglutarate (α-KG) dehydrogenase, and transketolase [39]. Therefore, we hypothesized that the mechanism by which thiamin primes plant defense is through changes in thiamin-dependent metabolic pathways, presuming that conversion of thiamin to ThDP is not rate-limiting, and that it leads to accumulation of higher ThDP cellular levels than normally observed. Because the direct response to thiamin treatment in distal/non-inoculated leaves yielded a limited transcriptional response, and following our observation that thiamin primes immune response with a limited treatment window (Fig. 1A), we surmised that the metabolomic response to foliar thiamin treatment may evolve earlier than the time course utilized in our RNA-seq experimental design, with thiamin treatment evoking changes in metabolite content in advance of changes in gene expression. Accordingly, we treated plants solely with 10 mM thiamin and surveyed the global metabolome via gas chromatography-mass spectrometry (GC-MS), sampling at 1, 6, and 12 hpt. A total of 92 analytes were identified in thiamin- and mock-treated samples (Fig. S6; Table S17), and significant upregulation and downregulation of these compounds by thiamin was determined via comparison to mock-treated samples at each time point.

At 1 hpt, we identified eight significantly upregulated compounds (Fig. 5A), including D-erythrose-4-phosphate, a direct product of thiamin-dependent transketolase activity in the Calvin cycle and the non-oxidative branch of the pentose phosphate pathway [40] (Fig. S7). At 1 hpt, we also detected in thiamin-treated samples significantly increased levels of glutamate, asparagine and putrescine, three metabolites that are synthesized from the TCA cycle intermediates α-KG and oxaloacetate, respectively (Fig. S7). Kyoto Encyclopedia of Genes and Genomes (KEGG) pathway enrichment analysis revealed that these eight analytes contribute to enrichment in three metabolic pathways: “alanine, aspartate, and glutamate metabolism”, via accumulation of asparagine and glutamate; and “glutathione metabolism” and “arginine and proline metabolism”, via accumulation of glutamate and putrescine (Table S18). At 6 hpt, five and 25 compounds significantly accumulated or decreased, respectively (Fig. 5B). Compounds that decreased contributed to six distinct metabolic pathways, corresponding to metabolism of various amino acids, galactose, and glyoxylate and dicarboxylate, as well as the citric acid cycle (Table S18, Fig. S7). At 12 hpt, only two upregulated compounds, sulfurol and 3,5- dihydroxyphenylglycine, were identified (Fig. 5C), with no corresponding significant pathway enrichment. Notably, across all time points, 4-methyl-5-thiazoleethanol (a.k.a. sulfurol) significantly accumulated as a result of thiamin treatment. Sulfurol is a thiazole moiety of thiamin, suggesting that all or a portion of the applied thiamin was rapidly degraded. However, GC-MS analysis revealed that the thiamin product (≥98%) applied to leaves contained traces of sulfurol, which could have contributed to the pool of sulfurol detected in leaf samples.

**Fig 5.**
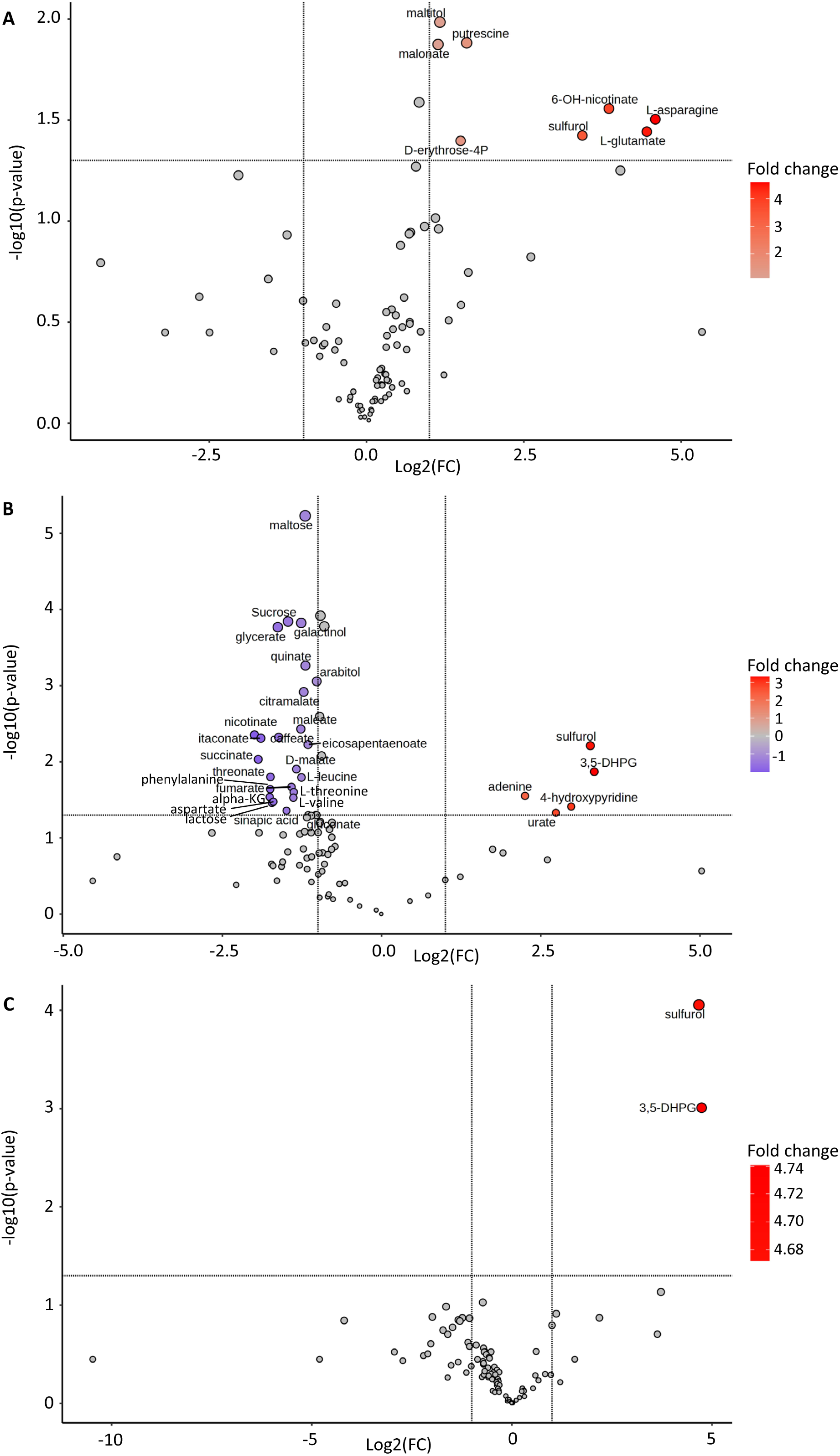
Volcano plots of metabolite concentrations in leaves of thiamin-treated versus mock- treated plants at 1 (A), 6 (B) and 12 (C) hpt. “FC”, fold change.

### Transcriptional responses to *A. solani* infection

To validate the transcriptional response to *A. solani*, we performed GO enrichment analysis utilizing the DEGs identified in “response to Alt (local)” and “response to Alt (thi)” (Fig. 6A-B; Tables S15-S16). Upregulated DEGs in both treatment groups were found to be significantly enriched for GO terms such as aromatic amino acid, terpene, phenylpropanoid, lignin, and chorismate biosynthesis, fatty acid oxidation, and response to oxidative stress. At 12 hpi, the “response to jasmonic acid” term was significantly enriched only in “response to Alt (thi)”, encompassing PR protein, lipoxygenase, JA-ZIM (JAZ) protein, ACC synthase, and ubiquitin ligase genes, as was the “response to fungus” term (Fig. 6A; Tables S10, S16). Additionally, genes corresponding to phenylalanine ammonia lyase (PAL) enzymes conferred significant enrichment of multiple PAL GO terms specifically in “response to Alt (thi)” at 12 hpi. At 48 hpi, “response to Alt (thi)” specifically exhibited significant enrichment for certain classes of GO terms, including: oxidative stress and lignin biosynthesis, via peroxidase genes; terpene biosynthesis; cell death, via phospholipase genes; and responses to chitin, via genes corresponding to U-box and LysM domain proteins.

**Fig 6.**
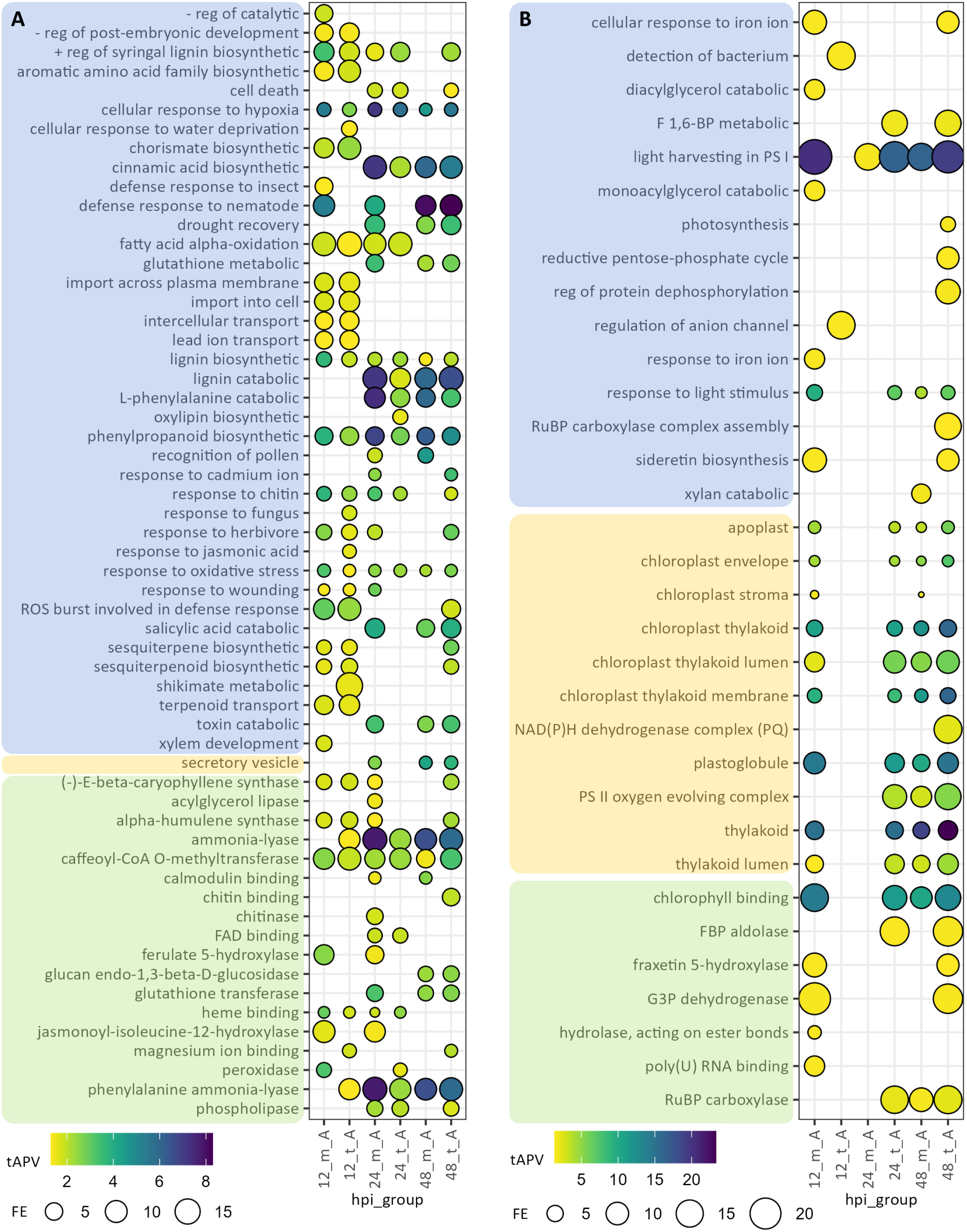
GO term enrichment of DEGs in “response to Alt (local)” (m_A) and “response to Alt (thi)” (t_A). DEGs were selected on the basis of |log_2_(fold-change)| ≥ 2 and adjusted p-value < 0.05. Enriched GO terms were selected on the basis of adjusted p-value (APV) < 0.05. “hpi”, hours post-inoculation. “tAPV”, -log_10_-transformed APV. “FE”, fold-enrichment. Blue-shaded terms, “Biological Process”; orange-shaded terms, “Cellular Component”; green-shaded terms, “Molecular Function”.

Analysis of GO enrichment among downregulated DEGs in “response to Alt (local)” and “response to Alt (thi)” highlighted a variety of GO terms associated with photosynthesis, such as “light harvesting in PSI”, “response to light stimulus”, “chlorophyll binding” activity, and cellular component terms associated with localization to the chloroplast (Fig. 6B; Tables S15- S16). Underlying genes comprise a range of gene products, such as chlorophyll-binding proteins, components of photosystems I and II, and Calvin cycle enzyme classes such as ribulose bisphoshate carboxylases (RuBisCo) and glyceraldehyde 3-phosphate (G3P) dehydrogenases (Tables S7-S12, S15-S16). Broadly, a pattern was observed wherein “response to Alt (local)” downregulated DEGs were enriched for photosynthesis-related functions and components by 12 hpi. However, thiamin treatment conferred a possible delay in this effect, with “response to Alt (thi)” downregulated DEGs exhibiting similar enrichments starting at 24 hpi, instead (Fig. 6B). Additionally, “response to Alt (thi)” downregulated DEGs were significantly enriched for multiple GO terms related to primary metabolism. At 48 hpi, downregulation of glycolytic fructose 1,6-bisphosphate (FBP) aldolase and G3P dehydrogenase genes yielded significant enrichment for GO terms corresponding to reductive pentose phosphate cycle and FBP processes and activities (Fig. 6B; Tables S12, S16). Interestingly, such downregulation was not observed in “response to Alt (local)”, indicating a direct effect of thiamin treatment on primary metabolic pathways.

## Discussion

Presently, control for biotic diseases of potato generally involves a combination of cultural and chemical methods, as well as the development of resistant potato varieties through breeding. Because such methods can be ineffective or expensive, or can carry detrimental environmental and health effects, it is important to evaluate alternative strategies to protect plants against microbial pathogens. Chemical immunity inducers are compounds that prime defense responses when applied to plant. In this study, we investigated thiamin, also known as vitamin B1, for its ability to induce immunity in potato to the fungal pathogen *A. solani*. Previous studies have tested the utility of thiamin treatments in several different plant-pathogen systems, but in potato, such work has thus far been limited to Potato Virus Y [28]. Furthermore, while aspects of thiamin-conferred disease resistance have been characterized through biochemical and histochemical assays [26, 41, 42], as well as genetically, via mutants and marker genes [23, 25, 42], the transcriptomics and metabolomics of immune priming by thiamin have not previously been investigated. Therefore, we sought to evaluate the effectiveness and potential modes of action of thiamin as a priming agent in potato, incorporating surveys of transcriptomic and metabolic responses via RNA-seq and GC-MS, respectively.

Utilizing detached leaf and whole plant assays, we observed that 10 mM was an optimal concentration for foliar applications of thiamin, yielding consistent reductions (32-52%) of foliar lesion area caused by the fungal pathogen *A. solani* (Table 1; Fig. 1B-C). This concentration is within the range of concentrations that have been reported in other phytopathosystems [23, 26]. We found that the protection provided by thiamin treatments was short lived, wherein protection against *A. solani* was abated when inoculations were performed later than 4 hpt (Fig. 1A). These results contrast with a previous study, in which thiamin conferred resistance to *Magnaporthe grisea* in rice, as well as *Pseudomonas syringae* in *Arabidopsis*, for up to 15 days after application [22]. This short-lived effect may be attributed to the rapid degradation of thiamin to its moieties, as indicated by the accumulation of 4-methyl-5-(2-hydroxyethyl)thiazole (sulfurol), the thiazole moiety of thiamin, shortly after foliar application (Fig. 5), although further quantification is needed to assess the contribution of the thiamin product to the detected sulfurol. Additionally, we observed that reapplication of thiamin one week post-initial treatment still conferred enhanced resistance (Fig. 1C), indicating that thiamin may be repeatedly applied to provide protection. However, additional application frequencies should be tested for strength and duration of immune priming.

We incorporated a bagged-leaf inoculation assay to evaluate the induction of systemic acquired resistance (SAR) by thiamin treatment and confirmed that thiamin immune priming does not act systemically (Fig. 1D). Conversely, previous studies utilizing different plant species observed thiamin-induced SAR responses after thiamin treatment. However, in these studies, SAR induction was indirectly measured, namely through monitoring of expression of SAR marker genes such as *PR-1*, instead of via direct disease assays [22, 26, 43]. Additionally, a study evaluating the activation of SAR by arachidonic acid in potato illustrated that arachidonic acid induced SAR against early and late blight, but only a local accumulation of SA, and no accumulation of PR-1 proteins in systemic tissues [44]. Via our gene expression analyses, we observed that application of thiamin induced expression of *PR-1* in non-inoculated plants at 24 hpi and enhanced the expression of *PR-1* in proximal tissue in response to *A. solani* at 48 hpi. Therefore, further evaluations of thiamin-induced SAR in potato may be augmented by incorporating additional experimental approaches, including alternative dosages and pathosystems, as well as broader surveys of SAR markers and putative mobile signals in proximal and distal tissues [45].

Utilizing RNA-seq and GO enrichment analysis to explore potential regulatory and response pathways activated by thiamin during immune priming, we observed thiamin to induce transcription of genes corresponding to chitinases, peroxidases, and protease inhibitors on distal, non-inoculated leaves (Fig. 3A; Tables S3-S4, S13). In response to *A. solani* infection, we found that thiamin treatment enhanced transcription of a variety of gene classes associated with pathogen defense, such as the biosynthesis of SA and phenylpropanoids, as indicated by enrichment for GO terms corresponding to shikimate biosynthesis and PAL activity, as well as peroxidases, PR proteins, U-box proteins, JAZ proteins, and chitinases, findings that are indicative of enhanced immune response, in comparison to mock-treated, *A. solani*-inoculated plants (Fig. 6; Tables S6-S11, S15-S16). Transcription of SA pathway, PR, and chitinase genes is well characterized in the context of plant-pathogen interactions, and a recent study reported activation of these gene classes, as well as the requirement of SA signaling, particularly in response to *A. solani* infections of potato [46]. The upregulation of genes involved in the biosynthesis of shikimate, a precursor of chorismate and substrate for SA and phenylalanine biosynthesis, uniquely in “response to Alt (thi)” at 12 hpi (i.e., without corresponding enrichment in “response to Alt (local)”) indicates that thiamin treatment may provide a boost towards defense responses in the first hours after inoculation. It is noteworthy that erythrose-4-phosphate, a precursor of shikimate produced by thiamin-dependent transketolases, was observed to accumulate 1 h post-thiamin treatment, absent pathogen challenge (Fig. 5A). Upon *A. solani* inoculation, higher thiamin availability may facilitate increased flux towards erythrose-4- phosphate, and subsequently shikimate biosynthesis, possibly necessitating increased expression of shikimate biosynthesis genes, suggesting a potential point of control of plant defense by thiamin.

Peroxidases have been characterized for diverse enzymatic functions within the cell, but are notably associated with the detoxification of ROS, which is generated during pathogen defense signaling and the cell death response; peroxidase genes have previously been observed to be upregulated in response to *A. solani* infections of potato [47, 48]. We also identified significant enrichment of genes, largely encoding phospholipase A2 proteins, involved in fatty acid metabolism at 48 hpi in “response to Alt (thi)” (Fig. 6A; Tables S12, S16). Furthermore, upregulated peroxidase- and fatty acid desaturase-encoding genes also contributed to enrichment of fatty acid metabolism functions in non-inoculated, thiamin-treated (“response to thi”) leaves at 48 hpi, and upregulated genes in these classes were identified at 12 and 24 hpi as well (Fig. 3; Tables S2-S4, S13). The phytohormone jasmonic acid (JA) is an integral part of defense against *A. solani* infections of potato, and fatty acids are precursors for JA (51, 52). Notably, in “response to thi” we observed upregulation of candidate JA biosynthesis genes encoding lipoxygenases (12 and 48 hpi) and an allene oxide synthase (48 hpi) (Tables S2, S4). Furthermore, we identified enrichment for the “response to jasmonic acid” GO term in “response to Alt (thi)” at 12 hpi (Fig. 6A; Table S16). These findings convey that, absent local pathogen infection, thiamin may prime immune response via activation of JA biosynthesis and signaling pathways, and upon infection, may act to activate these pathways at an earlier stage of the plant-pathogen interaction.

One of the most apparent consequences of thiamin treatment, in the context of infection with *A. solani*, is the temporal shift in the attenuation of photosynthesis-related gene expression. Photosynthesis metabolism can be highly modified by the immune response in plants, wherein crosstalk with defense phytohormones and ROS produces a shift in metabolic resources to mount a defense against pathogens [49, 50]. While GO enrichment analysis of downregulated DEGs indicated a broad reduction in expression of photosynthesis genes, such as chlorophyll-binding protein, photosystem (PS) component, and Calvin cycle enzyme genes, at 12 hpi, this effect was delayed in “response to Alt (thi)” leaves, with a similar pathway enrichment emerging instead at 24 hpi instead (Fig 6B; Tables S15-S16). Interestingly, thiamin also attenuated expression of chlorophyll-binding protein genes in non-inoculated (“response to thi”) leaves, conferring enrichment of photosynthesis GO terms, at 12 hpi (Fig. 3B; Tables S2, S13). In the *A. solani*- inoculated background (“response to thi (Alt)”), downregulation of chlorophyll and PS component and RuBisCo genes conferred GO term enrichments at 24 hpi (Fig. 4B; Tables S6, S14). Important defense phytohormones and signaling molecules like JA and nitric oxide are partially synthesized in the chloroplasts, and upon infection, transcriptional reprogramming is essential to produce these pro-defense molecules [51, 52]. Biosynthesis of fatty acids and aromatic amino acids is also compartmentalized to chloroplasts [53], and per our observations, expression of genes associated with metabolism of fatty acids and derivatives of aromatic amino acids is activated by pathogen infection and further enhanced by thiamin treatment. Accordingly, thiamin treatment is likely attenuating photosynthesis prior to infection with *A. solani*, priming defense activities against future infection, and/or enhancing immune response post-infection, possibly with an accompanying shift from photosynthesis to lipid and aromatic amino acid metabolism in the chloroplast.

Through “response to Alt (thi)” samples, we also observed thiamin to further suppress the expression of genes involved in primary metabolism, including G3P dehydrogenases and FBP aldolases, at 48 h-post *A. solani* infection (Fig. 6B; Tables S12, S16). Primary metabolic pathways such as glycolysis, the citric/tricarboxylic acid (TCA) cycle, and the pentose phosphate cycle are closely intertwined with photosynthesis, and ThDP is a known cofactor for key enzymes in these pathways, including transketolase, pyruvate dehydrogenase, and α-KG dehydrogenase [20]. Interestingly, three intermediates of the citric acid cycle, i.e., α-KG, fumarate and succinate, were depleted at 6 hpt compared to the control, suggesting an overall decrease in the flux of pyruvate towards the citric acid cycle, despite that two thiamin-dependent enzymes, pyruvate dehydrogenase and α-KG dehydrogenase, contribute to this pathway. Valine and leucine pools were also depleted at 6 hpt. In this case, increased activity of thiamin-dependent branched-chain α-ketoacid dehydrogenases, which catalyze the decarboxylation of the branched-chain α-ketoacids derived from valine and leucine (and isoleucine), may have contributed to increased degradation rate of valine and leucine, thereby decreasing their pools. Therefore, the priming of immune activity by thiamin may also result from a perturbation of primary metabolism that consequently shifts resources to activate pathogen defense pathways. Accordingly, future research should investigate thiamin-induced metabolic reprogramming in greater detail, focusing on the metabolite flux among the interconnected metabolic pathways that incorporate thiamin-dependent enzyme activities, with the goal of characterizing the resulting impacts to primary metabolism that inform the activation of immune priming by exogenous applications of thiamin.

## Conclusions

In conclusion, we have shown that foliar applications of thiamin decrease the size of lesions caused by the necrotrophic fungal pathogen *A. solani* in potato, indicating that thiamin could be included as part of an early blight management plan. However, additional field-scale research is warranted to test its efficacy in a commercial production environment. Through metabolites analyses, we have also shown that the mode of action of thiamin in priming plant defenses in the absence of *A. solani* involves a first phase where some metabolic intermediates of the TCA and Calvin cycles accumulate, followed by a second phase that seems to reverse course with decreased accumulation of some TCA cycle intermediates. Furthermore, transcriptomic analyses showed that this second phase is followed by repression of the expression of genes involved in photosynthesis by thiamin, and the increased expression of defense genes, such as JA-associated genes. Upon *A. solani* infection, thiamin delayed repression of the expression of photosynthesis-associated genes, further repressed the expression of genes involved in primary metabolism, and further enhanced the expression of genes associated with plant defenses.

## Electronic supplementary material

Table S1. Quality control metrics for RNA sequencing data.

Table S2. Differentially expressed genes (DEGs) in “response to thi”, 12 hours post-inoculation (hpi).

Table S3. Differentially expressed genes (DEGs) in “response to thi”, 24 hours post-inoculation (hpi).

Table S4. Differentially expressed genes (DEGs) in “response to thi”, 48 hours post-inoculation (hpi).

Table S5. Differentially expressed genes (DEGs) in “response to thi (Alt)”, 12 hours post-inoculation (hpi).

Table S6. Differentially expressed genes (DEGs) in “response to thi (Alt)”, 24 hours post-inoculation (hpi).

Table S7. Differentially expressed genes (DEGs) in “response to Alt (local)”, 12 hours post-inoculation (hpi).

Table S8. Differentially expressed genes (DEGs) in “response to Alt (local)”, 24 hours post-inoculation (hpi).

Table S9. Differentially expressed genes (DEGs) in “response to Alt (local)”, 48 hours post- inoculation (hpi).

Table S10. Differentially expressed genes (DEGs) in “response to Alt (thi)”, 12 hours post- inoculation (hpi).

Table S11. Differentially expressed genes (DEGs) in “response to Alt (thi)”, 24 hours post- inoculation (hpi).

Table S12. Differentially expressed genes (DEGs) in “response to Alt (thi)”, 48 hours post- inoculation (hpi).

Table S13. Enriched Gene Ontology (GO) terms identified in differentially expressed genes (DEGs) in “response to thi.”

Table S14. Enriched Gene Ontology (GO) terms identified in differentially expressed genes (DEGs) in “response to thi (Alt).”

Table S15. Enriched Gene Ontology (GO) terms identified in differentially expressed genes (DEGs) in “response to Alt (local).”

Table S16. Enriched GO terms identified in DEGs in “response to Alt (thi).”

Table S17. Metabolite concentrations in leaf samples in thiamin- or mock-treated plants at 1, 6 and 12 hpt as determined by GC-MS.

Table S18. Pathway enrichment of metabolomics data following thiamin treatment.

Figure S1. Schema for treatment groups (left) and comparison groups (right) for RNA-seq differential expression analysis.

Figure S2. Pictures of lesions at 12, 24 and 48 hpi with *Alternaria solani*. Figure S3. qRT-PCR gene expression analysis of *PR-1*.

Figure S4. Principal Component Analysis (PCA) plot of RNA-seq samples.

Figure S5. Volcano plots for differentially expressed genes (DEGs) at 12 (left), 24 (center), and 48 (right) hours post-inoculation.

Figure S6. Heatmap of concentrations of metabolites analyzed by GC-MS. M1, M6 and M12: mock-treated plants at 1, 6 and 12 hpt; T1, T6 and T12: thiamin-treated plants at 1, 6 and 12 hpt. Data represent averages of 4 biological replicates.

Figure S7. Simplified schema of the Calvin cycle, glycolysis, the TCA cycle, and α-ketoacids catabolism with thiamin-dependent enzymatic steps. In orange squares are thiamin-dependent enzymes. TK, transketolase; PDH, pyruvate dehydrogenase; KGDH, 2-oxoglutarate (α-KG) dehydrogenase; BCKDH, branched-chain amino acids ketodehydrogenase. A. Thiamin-dependent pathways where metabolites accumulated at 1 hpt with thiamin (in red text). B. Thiamin-dependent pathways where metabolites decreased at 6 hpt with thiamin (in blue text).

## Declarations

### Availability of data and materials

Raw Illumina sequencing reads are deposited at the NCBI Sequence Read Archive under the BioProject accession number PRJNA1149021.

### Competing interests

The authors declare that they have no competing interests.

### Funding

Trenton Berrian was supported by grants from the Western Sustainable Agriculture Research and Education (Award No. GW22-239) and the USDA National Institute of Food and Agriculture (Award No. 2021-38420-34064). This project was also supported by a grant from the USDA-ARS/State Partnership program to CRC and AJG.

### Authors’ contributions

T.B., M.F., C.C. and A.G. designed the experiments. T.B. collected the samples. T.B. and M.F. performed the laboratory experiments. C.R. and J.A. contributed the GC-MS analyses. T.B., M.F., C.C. and A.G. analyzed the data. T.B., M.F. and A.G. wrote the manuscript. C.C. revised and edited the manuscript. C.C. and A.G. acquired the funding and supervised the project. All authors read and approved the final manuscript.

## Acknowledgments

We would like to thank Dr. Barry Pryor (University of Arizona) for donating *Alternaria solani* isolate used in this study.

## Notes

### Competing Interest Statement

The authors have declared no competing interest.

