## Supplementary Figures for "Thiamin priming to control early blight in potato: investigation of its effectiveness and molecular mechanisms": Supplementary Figures.pdf

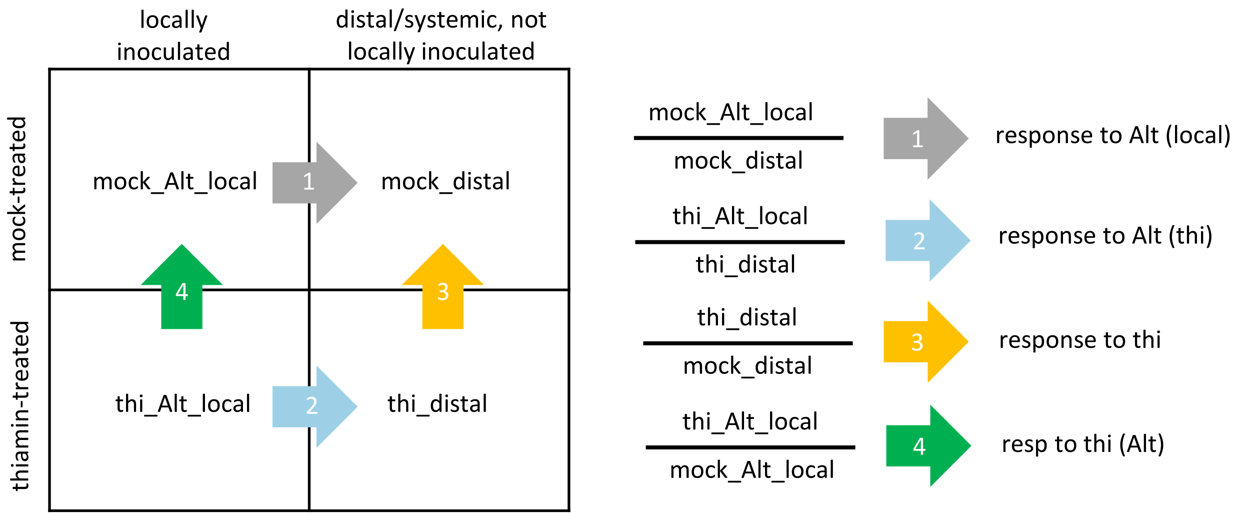


**Figure S1.** Schema for treatment groups (left) and comparison groups (right) for RNA-seq differential expression analysis.

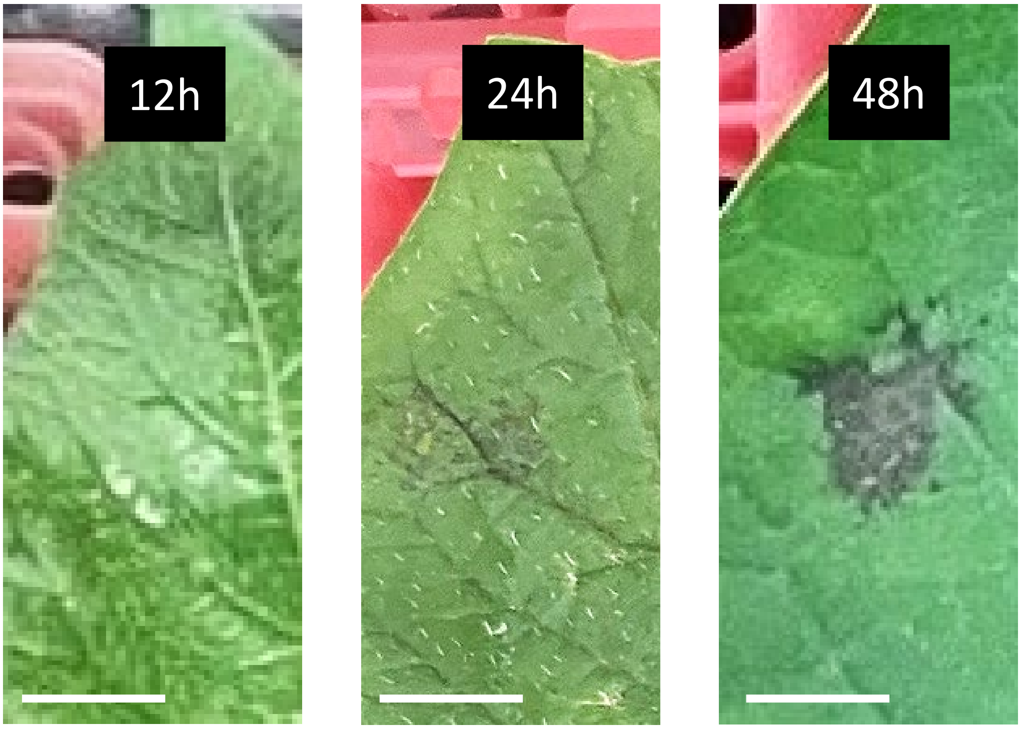


**Figure S2.** Pictures of lesions at 12, 24 and 48 hpi with *Alternaria solani*.

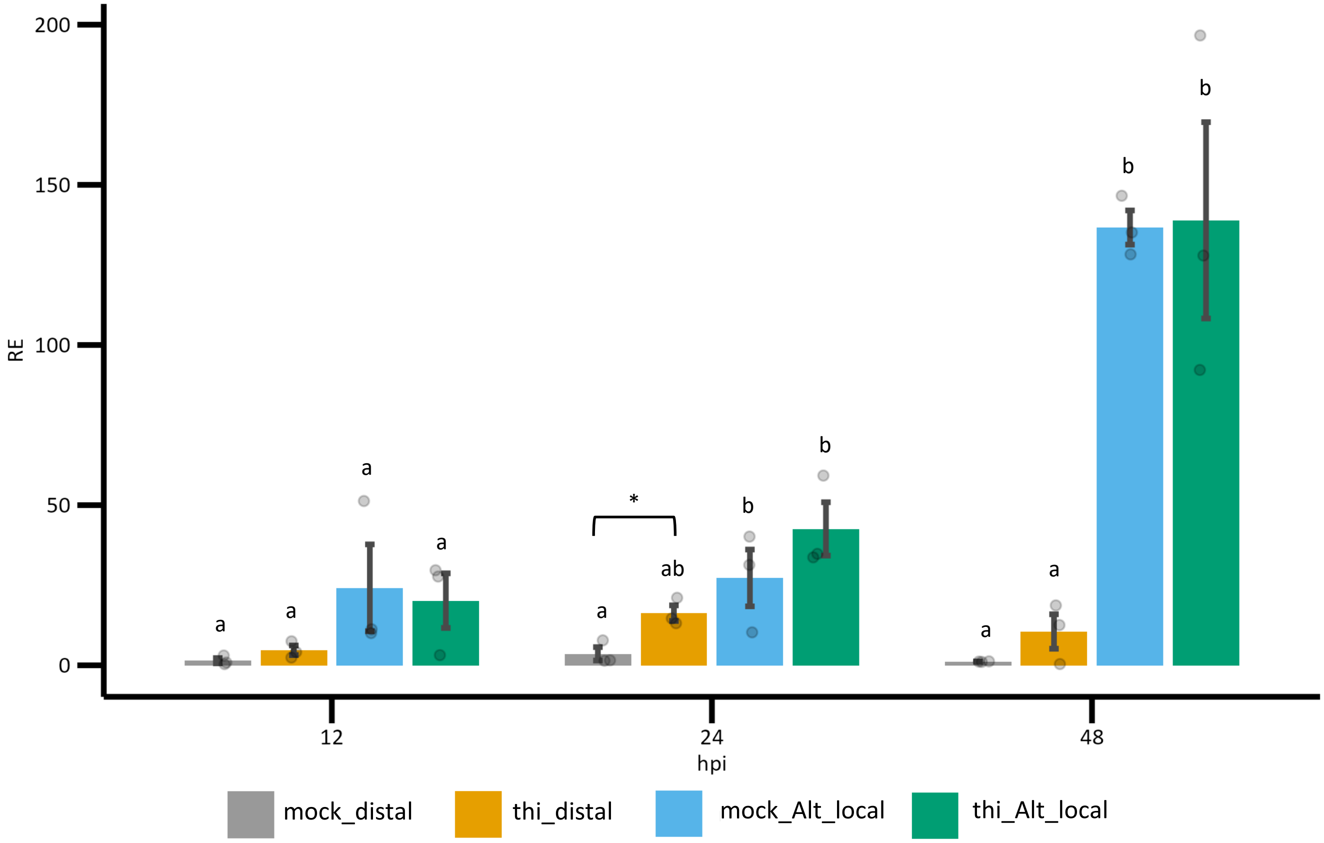


**Figure S3.** qRT-PCR gene expression analysis of *PR-1*. "RE", relative expression. Letters indicate statistical significance classes determined via ANOVA with post hoc Tukey's test. Asterisk indicates statistical significance ($p < 0.05$) determined via 1-way T-test.

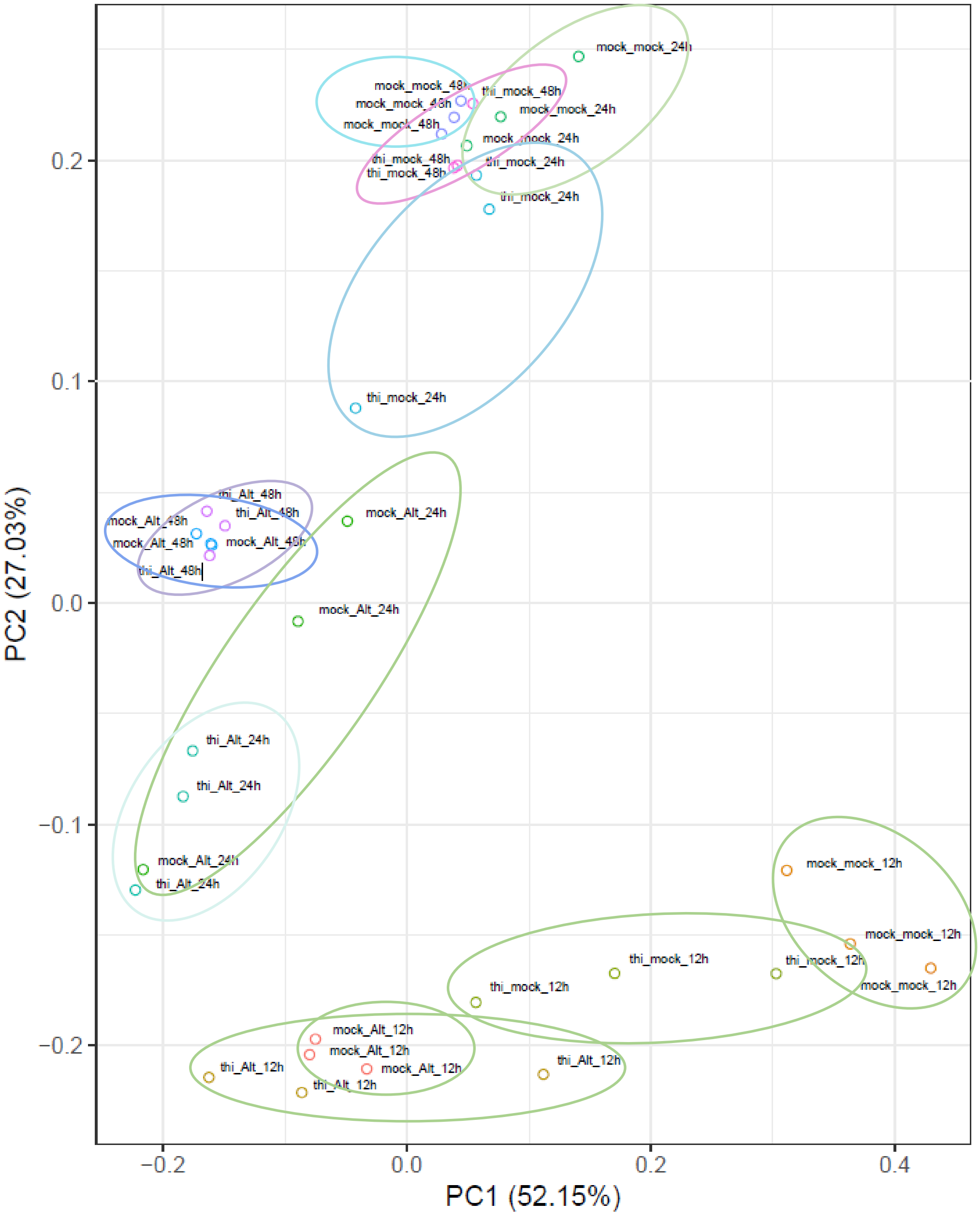


**Figure S4.** Principal Component Analysis (PCA) plot of RNA-seq samples.

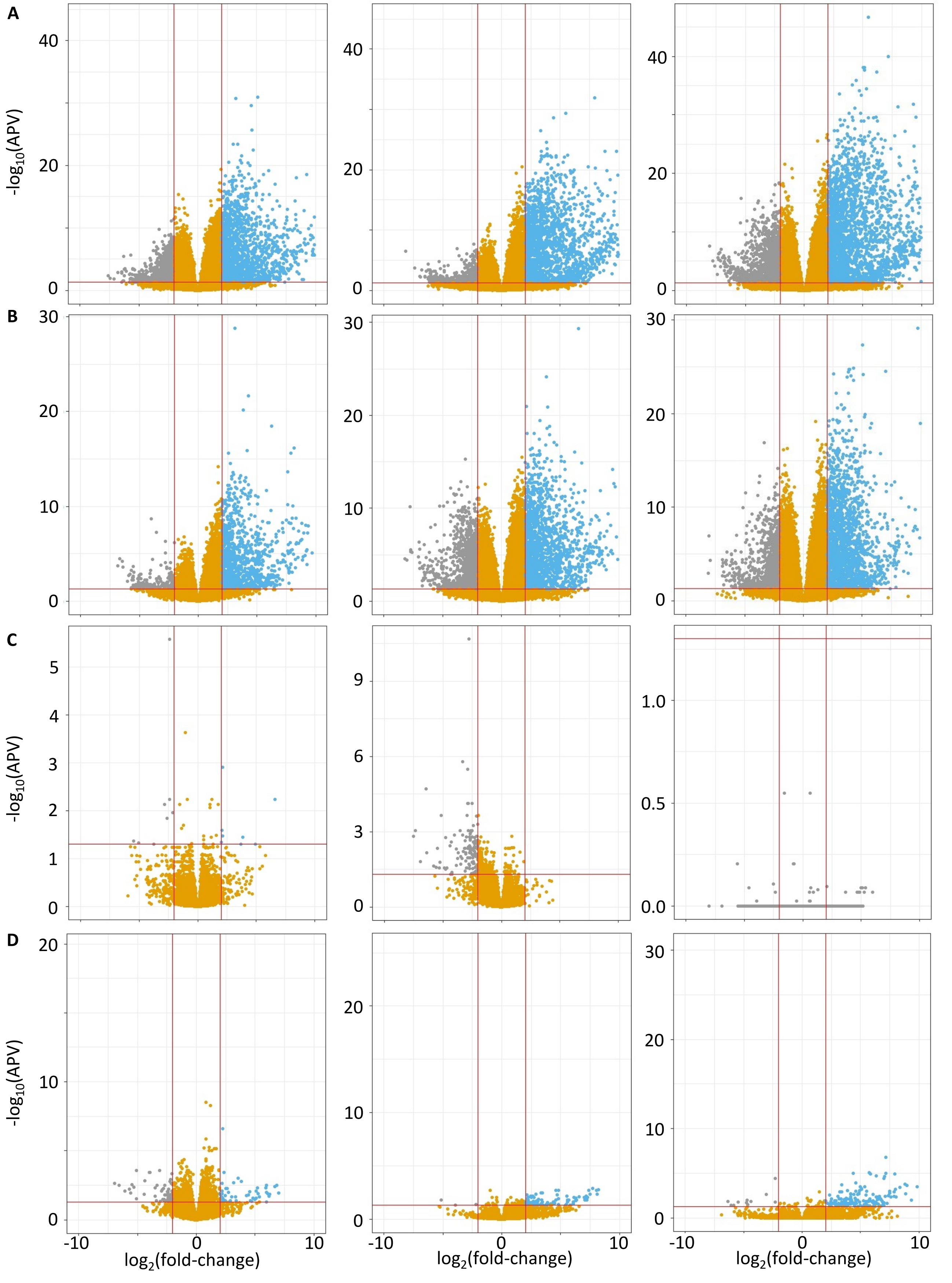


**Figure S5.** Volcano plots for differentially expressed genes (DEGs) at 12 (left), 24 (center), and 48 (right) hours post-inoculation. (A), “response to Alt (local).” (B), “response to Alt (thi).” (C), “response to thi (Alt).” (D), “response to thi.” “APV”, adjusted p-value.

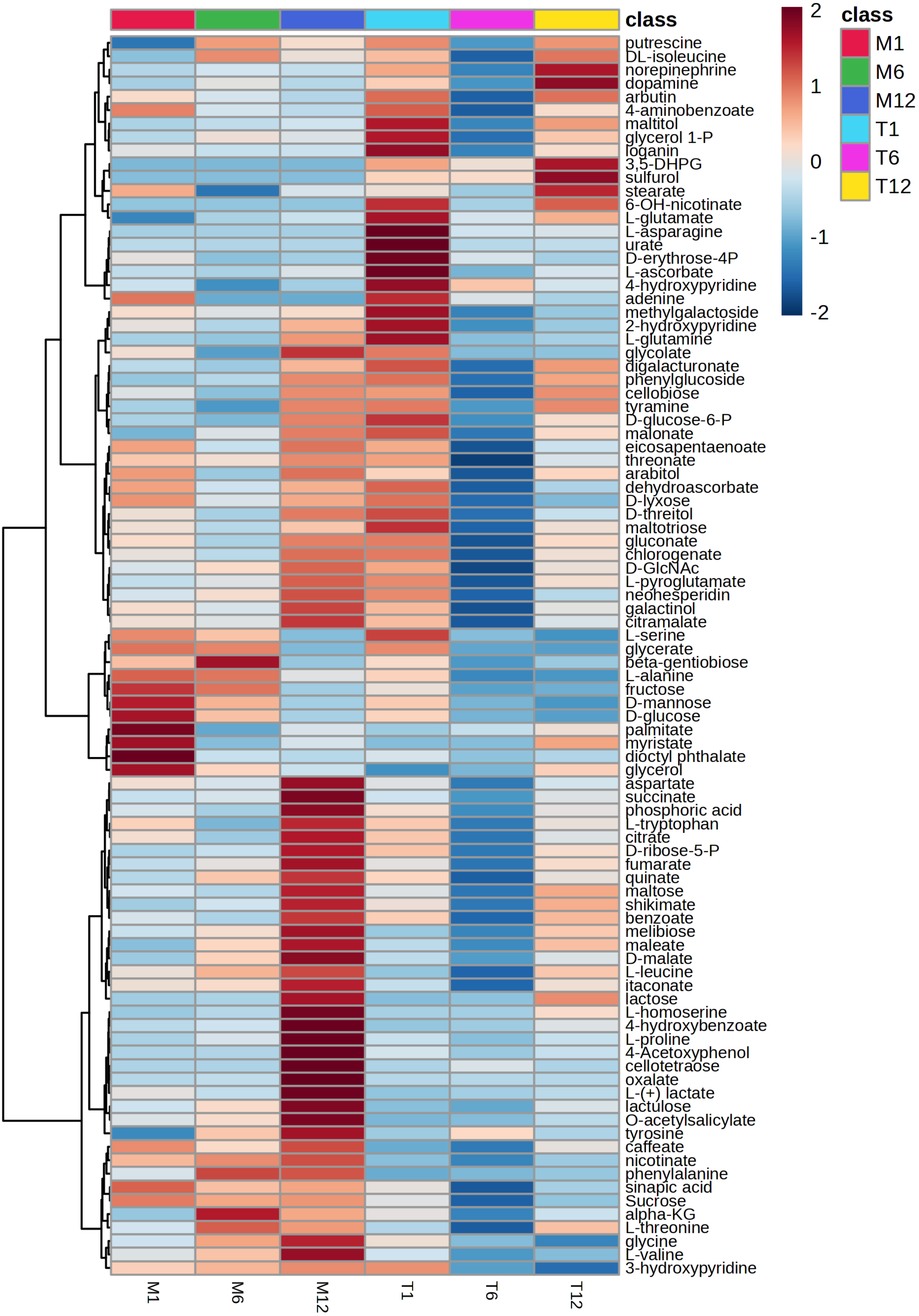


**Figure S6.** Heatmap of concentrations of metabolites analyzed by GC-MS. M1, M6 and M12: mock-treated plants at 1, 6 and 12 hpt; T1, T6 and T12: thiamin-treated plants at 1, 6 and 12 hpt. Data represent averages of 4 biological replicates.

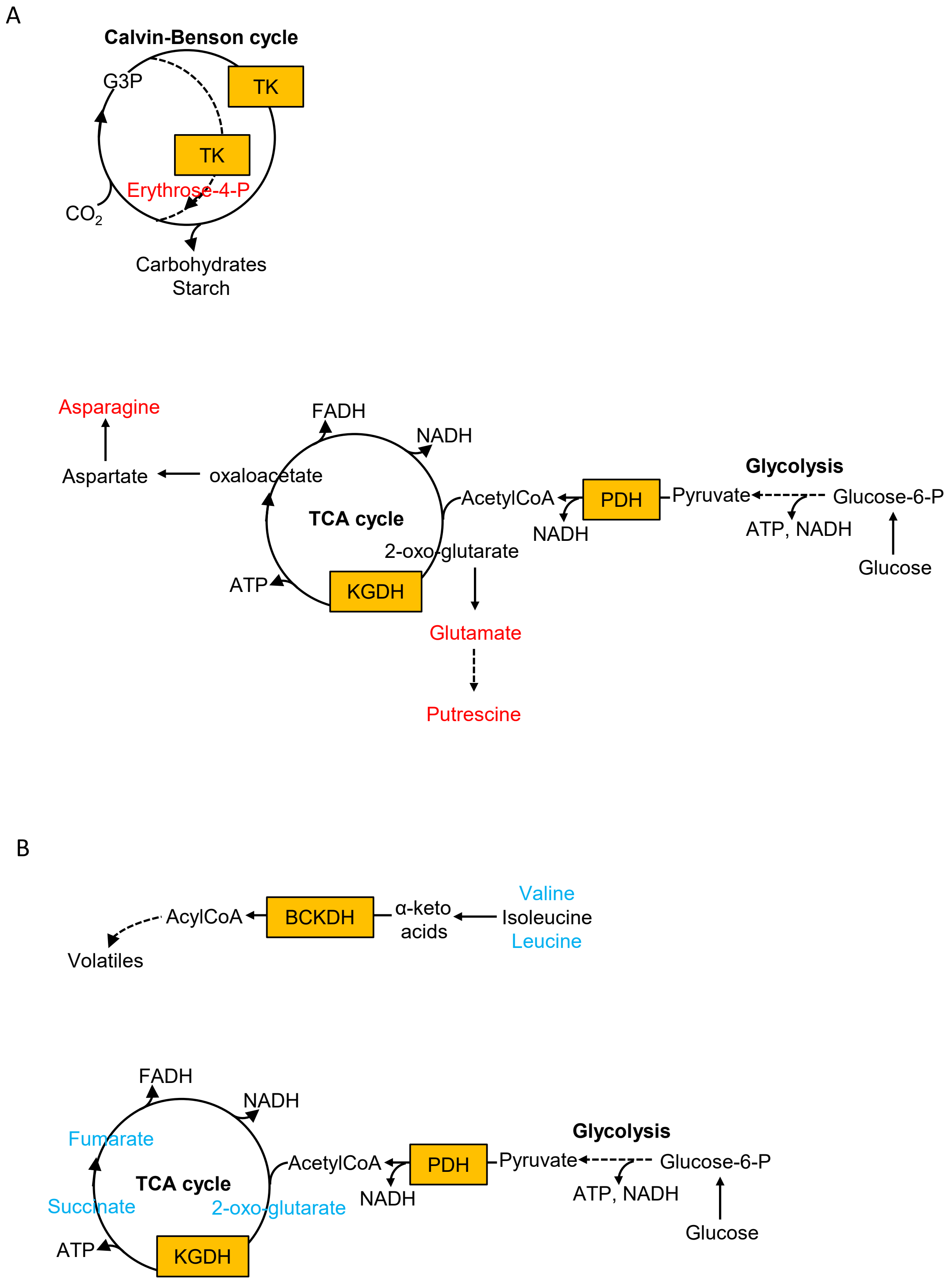


**Figure S7.** Simplified schema of the Calvin cycle, glycolysis, the TCA cycle, and α-ketoacids catabolism with thiamin-dependent enzymatic steps. In orange squares are thiamin-dependent enzymes. TK, transketolase; PDH, pyruvate dehydrogenase; KGDH, 2-oxoglutarate (α-KG) dehydrogenase; BCKDH, branched-chain amino acids ketodehydrogenase. A. Thiamin-dependent pathways where metabolites accumulated at 1 hpt with thiamin (in red text). B. Thiamin-dependent pathways where metabolites decreased at 6 hpt with thiamin (in blue text).
